# Genomic analyses identify biological processes and functions between M1 and M2 macrophages

**DOI:** 10.1101/2022.02.28.482314

**Authors:** Tingxiang Chang, Tingting Zhang, Hanming Gu, Jing Wang

## Abstract

Macrophages can be induced by a variety of factors to change their phenotype and functions. M1 and M2 macrophages play converse roles during inflammation or other diseases. However, the mechanism and function of M1 and M2 macrophages are still not clear. This study aims to identify the different molecules and functions by analyzing the RNA-seq data. The GSE189354 was created by the Illumina HiSeq 4000 (Mus musculus). The KEGG and GO analyses indicated the Phagosome and Hematopoietic cell lineage are the major biological processes between M1 and M2 macrophages. Moreover, we further identified ten molecules including Il6, Il1b, Tlr2, Myc, Fn1, Itgax, Cxcl10, Ccl5, Cxcr4, and Pparg. Therefore, our study may provide novel knowledge of innate immunity.

## Introduction

Macrophages have diverse features in maintaining an organism’s integrity by either triggering the inflammatory responses or repairing tissues under specific conditions^1^. There are different phenotypes of macrophages and each one has its own characteristics and functions^2^.

M1 macrophages produce pro-inflammatory cytokines and present strong microbicidal properties^3^. The classical stimuli include IFN-γ, lipopolysaccharide (LPS), and granulocyte-macrophage colony-stimulating factor (GM-CSF)^4^. M1 macrophages are characterized by an increased level of cytokines such as IL1B, IL12, and IL18^5^. M2 macrophages are activated through non-classical pathways and the stimuli of M2 macrophages are CSF-1, IL4, IL10, TGFβ, and IL13^6^. M2 macrophages are related to parasites, tissue remodeling, and angiogenesis^7^. The arginase 1 (Arg1), MMR (Mrc1), resistin-like molecule α (FIZZ1), and chitinase-like protein Ym1 are considered M2 macrophages marker genes^1^. M1 macrophages are involved in controlling autoimmune diseases such as inflammatory bowel disease (IBD), multiple sclerosis (MS), rheumatoid arthritis (RA), and Type 1 diabetes. M2 macrophages are involved in allergic asthma and parasite infection^8^.

In this study, we determined the signaling pathways of M1 and M2 macrophages by using the RNA-seq data. We identified a number of DEGs and biological processes and constructed the protein-protein interaction (PPI) network. The DEGs and PPI network may provide deep insights into macrophage functions.

## Methods

### Data resources

Gene dataset GSE189354 was collected from the GEO database. The data was produced by the Illumina HiSeq 4000 (Mus musculus) (University of Michigan, 2800 Plymouth Rd, Ann Arbor, MI, USA). The analyzed dataset includes muscles treated by 3 groups of M1 macrophages and 3 groups of M2 macrophages.

### Data acquisition and processing

The data were conducted by the R package as previously described^9-17^. We used a classical t-test to identify DEGs with P<0.01 and fold change ≥1.5 as being statistically significant.

The Kyoto Encyclopedia of Genes and Genomes (KEGG) and Gene Ontology (GO) KEGG and GO analyses were performed by the R package (clusterProfiler) and Reactome. P<0.05 was considered statistically significant.

### Protein-protein interaction (PPI) networks

The Molecular Complex Detection (MCODE) was used to create the PPI networks. The significant modules were produced from constructed PPI networks and String networks. The signaling pathway analyses were performed by using Reactome (https://reactome.org/), and P<0.05 was considered significant.

## Results

### Identification of DEGs between M1 and M2 macrophages

To determine the genetic difference between M1 and M2 macrophages, we analyzed the RNA-seq data from LPS-induced M1 and IL4 induced-M2 macrophages. A total of 624 genes were identified with the threshold of P < 0.01. The top up- and down-regulated genes between M1 and M2 macrophages were identified by the heatmap and volcano plot (Figure 1). The top ten DEGs were listed in Table 1.

**Table 1.**

| **Entrez gene** | **Gene symbol** | **Fold-change** | **Regulation** |
| --- | --- | --- | --- |
| Top 10 down-regulated DEGs |  |  |  |
| 149233 | Il23r | -8.951305238 | Down |
| 952 | Cd38 | -8.38970171 | Down |
| 2357 | Fpr1 | -8.309375697 | Down |
| 20210 | Saa3 | -8.301682841 | Down |
| 56241 | Susd2 | -7.901408648 | Down |
| 342510 | Cd300e | -7.883115517 | Down |
| 6590 | Slpi | -7.840214618 | Down |
| 6352 | Ccl5 | -7.359987709 | Down |
| 629 | Cfb | -7.304566533 | Down |
| 149233 | Il23r | -8.951305238 | Down |
| Top 10 up-regulated DEGs |  |  |  |
| 57262 | Retnla | 9.835959115 | up |
| 93726 | Rnase2a | 9.342942891 | up |
| 216864 | Mgl2 | 9.179234342 | up |
| 55806 | Hr | 8.526610924 | up |
| 6369 | Ccl24 | 8.108896497 | up |
| 2615 | Lrrc32 | 7.827037348 | up |
| 3687 | Itgax | 7.641890704 | up |
| 128506 | Ocstamp | 7.471849941 | up |
| 245972 | Atp6v0d2 | 7.432008132 | up |
| 100043726 | Gm42835 | 9.835959115 | up |

**Figure 1.**
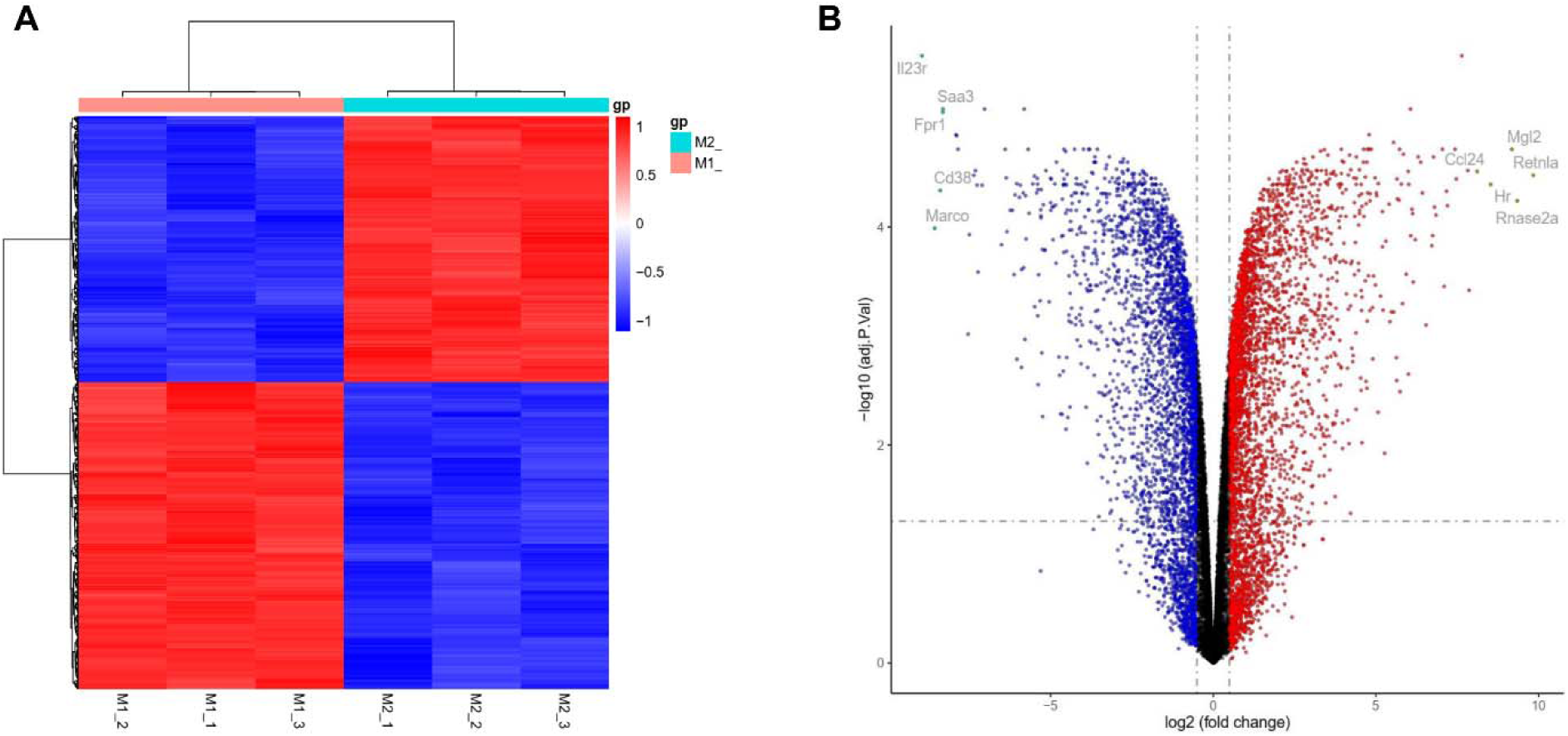
Heatmap and volcano plot were created between M1 and M2 macrophages. (A) Heatmap of significant DEGs. Significant DEGs (P < 0.01) were used to produce the heatmap. (B) Volcano plot for DEGs between M1 and M2 macrophages. The most significantly changed genes are highlighted by grey dots.

### Enrichment analysis of DEGs between M1 and M2 macrophages

To identify the biological processes between M1 and M2 macrophages, we performed the KEGG and GO analyses (Figure 2). We identified the top ten KEGG items including “Phagosome”, “Hematopoietic cell lineage”, “Tuberculosis”, “Graft−versus−host disease”, “Inflammatory bowel disease”, “Type I diabetes mellitus”, “Rheumatoid arthritis”, “Antigen processing and presentation”, “Leishmaniasis”, and “Allograft rejection”. We figured out the top ten biological processes (BP) of GO, including “adaptive immune response based on somatic recombination of immune receptors built from immunoglobulin superfamily domains”, “positive regulation of response to external stimulus”, “cytokine−mediated signaling pathway”, “leukocyte cell−cell adhesion”, “leukocyte migration”, “regulation of leukocyte cell−cell adhesion”, “regulation of response to biotic stimulus”, “negative regulation of cytokine production”, “response to interferon−gamma”, and “antigen processing and presentation”. We identified the cellular components (CC) of GO, including “apical part of cell”, “receptor complex”, “late endosome”, “endosome membrane”, “endocytic vesicle”, “plasma membrane signaling receptor complex”, “phagocytic vesicle”, “MHC protein complex”, “phagocytic vesicle membrane”, and “MHC class II protein complex”. We also identified the top ten molecular functions of GO, including “peptide binding”, “amide binding”, “cytokine receptor binding”, “carbohydrate binding”, “immune receptor activity”, “cytokine binding”, “peptide antigen binding”, “T cell receptor binding”, “CD8 receptor binding”, and “MHC class II protein complex binding”.

**Figure 2.**
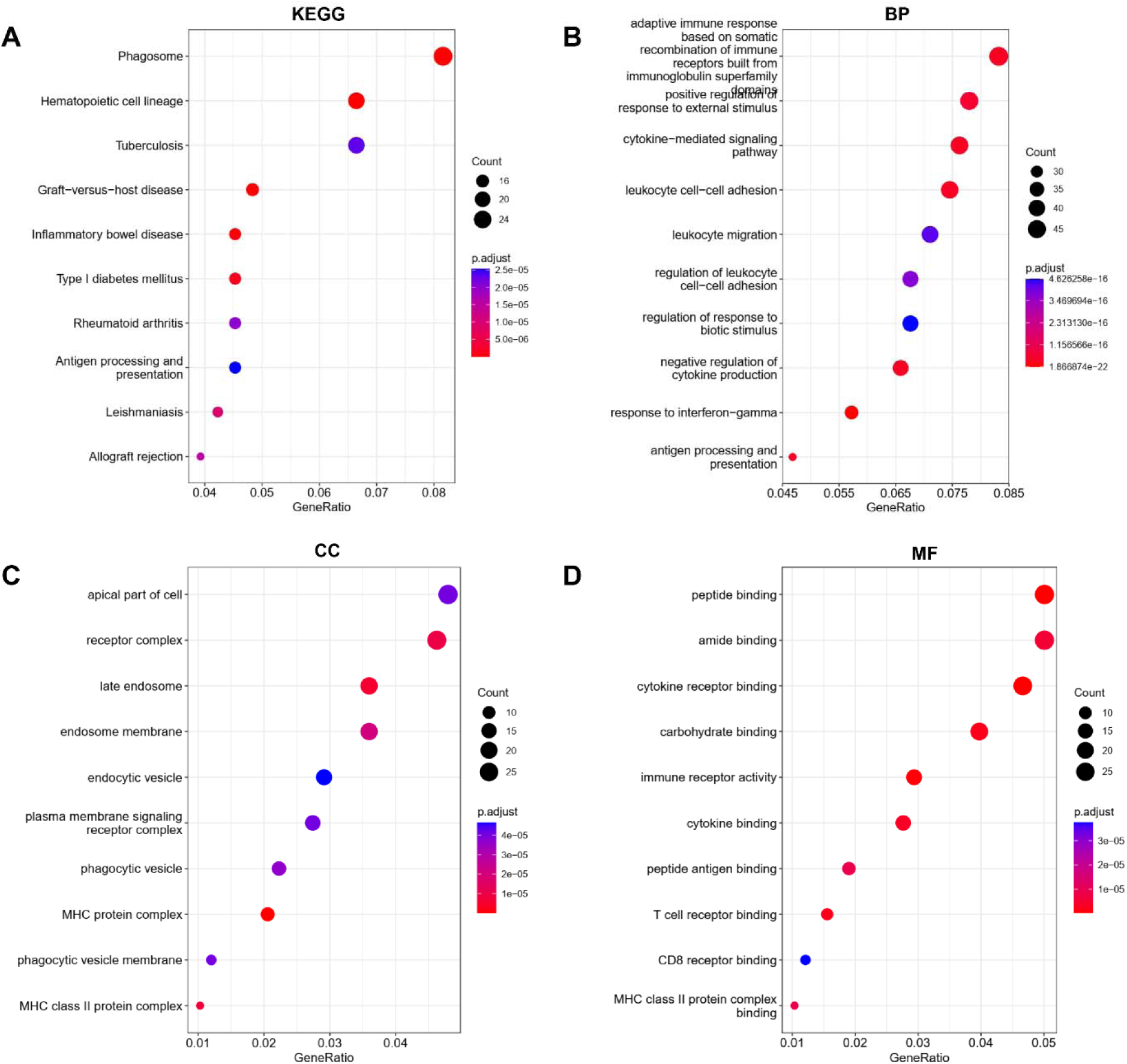
KEGG and GO analyses of DEGs between M1 and M2 macrophages. (A) KEGG analysis, (B) Biological processes, (C) Cellular components, (D) Molecular functions.

### PPI network between M1 and M2 macrophages

To further study the potential relationships between the DEGs, we created the PPI network by using the Cytoscape. The combined score > 0.2 was used as a cutoff to construct the PPI network by linking 510 nodes and 3549 edges. Table 2 indicated the top ten genes with the highest degree scores. The top two clusters were shown in Figure 3. We then evaluated the DEGs and PPI network by Reactome map (Figure 4) and then identified the top ten significant biological processes including “Signaling by Interleukins”, “Interleukin-4 and Interleukin-13 signaling”, “Cytokine Signaling in Immune system”, “Immune System”, “Neutrophil degranulation”, “Interleukin-10 signaling”, “Interferon Signaling”, “Interleukin-35 Signalling”, “Transcriptional regulation of white adipocyte differentiation”, and “RUNX3 Regulates Immune Response and Cell Migration” (Supplemental Table S1).

**Table 2.** Top ten genes demonstrated by connectivity degree in the PPI network.

| **Gene symbol** | **Gene title** | **Degree** |
| --- | --- | --- |
| Il6 | Interleukin 6 | 122 |
| Il1b | Interleukin 1 beta | 119 |
| Tlr2 | Toll like receptor 2 | 78 |
| Myc | MYC proto-oncogene | 72 |
| Fn1 | Fibronectin 1 | 70 |
| Itgax | Integrin subunit alpha X | 62 |
| Cxcl10 | C-X-C motif chemokine ligand 10 | 62 |
| Ccl5 | C-C motif chemokine ligand 5 | 61 |
| Cxcr4 | C-X-C motif chemokine receptor 4 | 61 |
| Pparg | Peroxisome proliferator activated receptor gamma | 50 |

**Figure 3.**
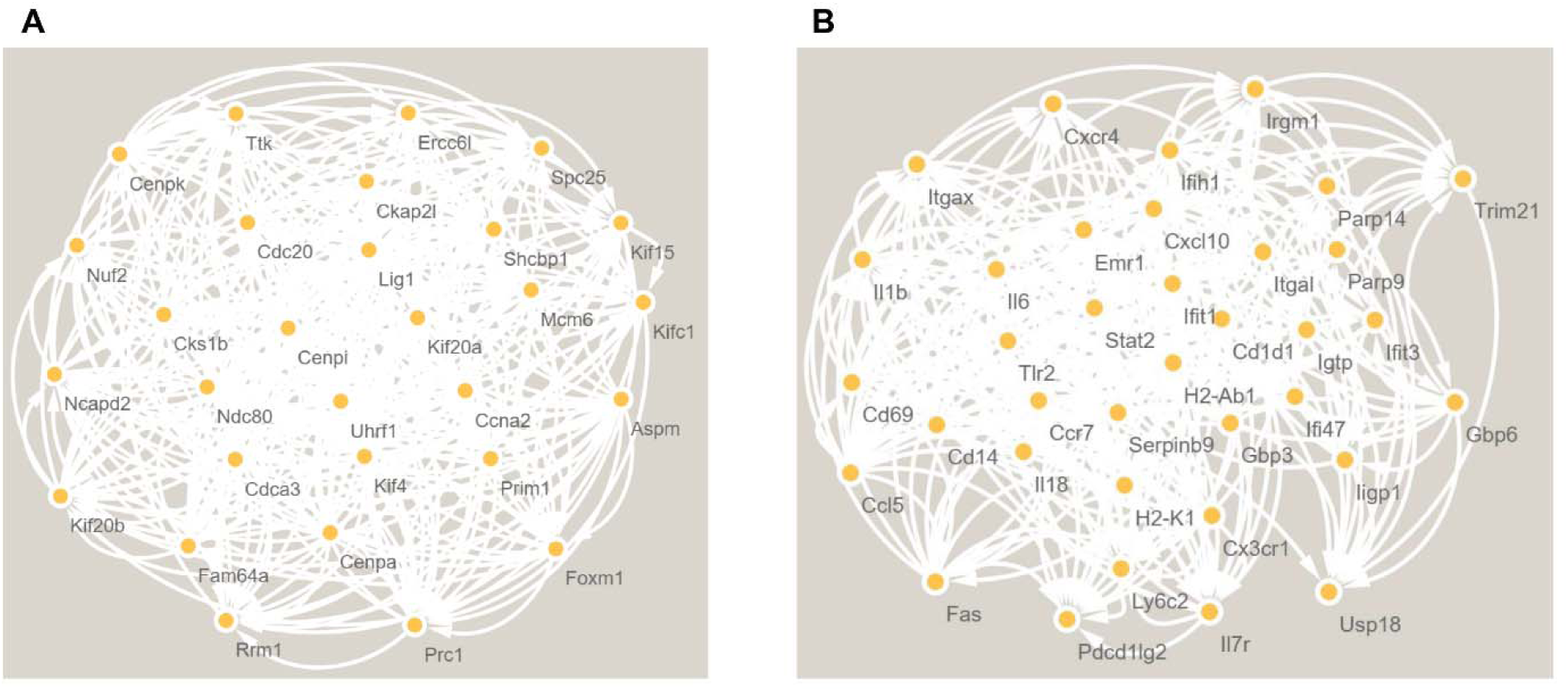
The PPI network analyses of DEGs between M1 and M2 macrophages. The cluster (A) and cluster (B) were constructed by MCODE.

**Figure 4.**
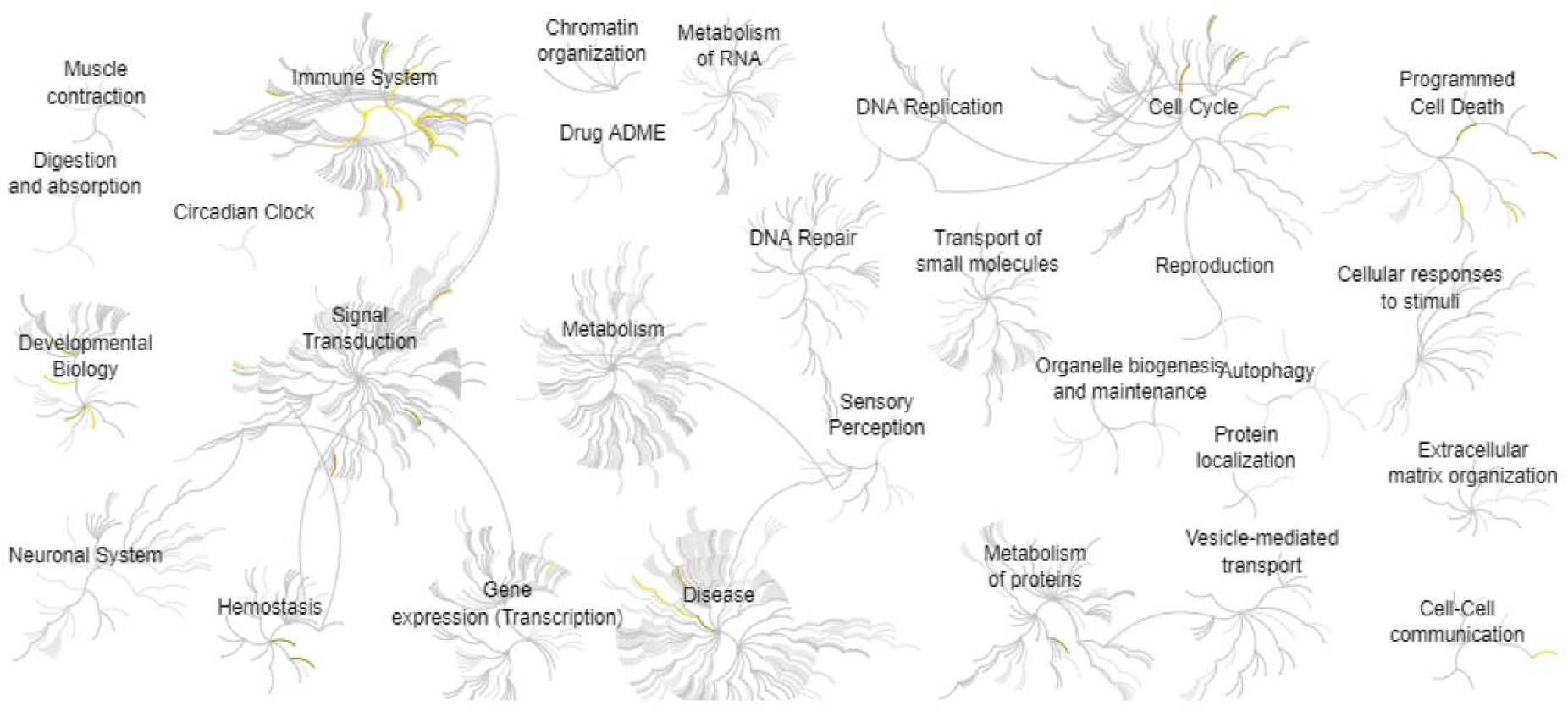
Reactome map representation of the significant biological processes between M1 and M2 macrophages.

## Discussion

Macrophages can be regulated by various factors to alter their phenotype and functions^18^. M1 macrophages are capable of proinflammatory responses and producing cytokines. Conversely, M2 macrophages indicate the anti-inflammatory responses and repair damaged tissue^19^. In infected tissues, macrophages are polarized to M1 phenotype to prevent the pathogens, then the M1 macrophages are changed to M2 phenotype to repair damaged tissue^3^. Here, our study aims to elaborate on the molecular mechanism between M1 and M2 macrophages.

By analyzing the KEGG and GO data, we found that Phagosome and Hematopoietic cell lineage are the major biological processes between M1 and M2 macrophages. The study by Johnathan Canton et al found that M1 macrophages had a similar buffering power and proton leakage permeability but decreased proton-pumping activity compared with M2 phagosomes^20^. Ujjaldeep Jaggi et al demonstrated that the overexpression of M2 macrophages was related to increased phagocytosis and anti-inflammatory cytokines^21^. The study by Xuan Huang et al found Macrophages are a heterogeneous population of innate myeloid cells that are located in the hematopoietic system. M1 macrophage can be induced by IFNγ and LPS and M2 macrophages can be induced by several stimuli including IL4, IL13, immune complexes, glucocorticoids, IL10, and IL6^22^.

In this study, we also identified the association molecules by using the PPI network. Liliana M.Sanmarco et al found IL6 drives M2 macrophage polarization by mediating purinergic signaling and controls the lethal release of nitric oxide during Trypanosoma cruzi infection^23^. Circadian clocks and their downstream targets control a variety of biological processes including immunity, metabolism, aging, and cancer^24-34^. Kandis L. Adams et al found that LPS-induced IL-6 release was regulated in a circadian manner, peaking during subjective day. This rhythm was associated with daily variation in immune cell responsiveness^35^. IL1 and IL4 induced macrophages contain opposite effects on adipogenesis in humans^36^. Lilian Quero et al demonstrate that TLR2 inhibits the anti-inflammatory function of M2-like macrophages, indicating a chimeric M1/M2 phenotype^37^. MYC is a transcription factor involved in cell proliferation, survival, and angiogenesis. In M2 macrophage polarization, MYC is induced by IL4 and controls the expression of M2 genes^38^. GPCR and RGS regulate a couple of molecules to further the cell functions such as proliferation, survival, differentiation, and immunity^39-49^. The knockdown of GPCR protein TDAG8 reduces the c-Myc expression^50^. FN1 is a known M2 macrophage marker that can be induced by IL4^51^. ITGAX is a proinflammatory gene that is expressed in muscle and correlates positively with glycemia^52^. Veethika Pandey et al found CXCL10 creates an inflammatory microenvironment and inhibits the progression of pancreatic precancerous lesions^53^. Yuan Zhuang et al demonstrated that inhibition of CCL5-CCR5 enhances M1 macrophage polarization^54^. CXCL12-CXCR4 signaling promotes M1 macrophage accumulation and blocking the signaling inhibits the secretion of pro-inflammatory cytokines^55^. Qinyu Yao et al found PPARγ induces the integrin αVβ5 to enhance M2 macrophage polarization^56^.

In conclusion, our study identified the genetic difference between M1 and M2 macrophages. Phagosome and Hematopoietic cell lineage are the main biological pathways that were affected between M1 and M2 macrophages. Therefore, our study may provide insights into innate immunity.

## Author Contributions

Tingxiang Chang and Tingting Zhang: Methodology. Hanming Gu and Jing Wang: Conceptualization, Writing and Editing.

